# Neuronal responses to focused ultrasound are gated by pre-stimulation brain rhythms

**DOI:** 10.1101/2021.08.06.455443

**Authors:** Duc Nguyen, Elisa Konofagou, Jacek P. Dmochowski

## Abstract

**Background:** Owing to its high spatial resolution and penetration depth, transcranial focused ultrasound stimulation (tFUS) is one of the most promising approaches to non-invasive neuromodulation. Identifying the impact of the stimulation waveform and endogenous neural activity on neuromodulation outcome is critical to harnessing the potential of tFUS.

**Objective:** Here we tested a new form of tFUS where the amplitude of the ultrasonic waveform is modulated at a rate much slower than the operating frequency. Moreover, we sought to identify the relationship between pre-stimulation neural activity and the neuronal response to tFUS.

**Methods:** We applied three minutes of amplitude modulated (AM) tFUS at 40 Hz to the rat hippocampus while recording local field potentials (LFP) and multi-unit activity (MUA)from the sonicated region. To assess the role of AM, we also tested continuous-wave (CW) stimulation.

**Results:** AM tFUS reduced firing rate during and immediately after stimulation. On the other hand, CW tFUS produced an acute firing rate increase that was abolished after sonication. For both waveforms, firing rate changes were stronger in units exhibiting high baseline LFP power, particularly in the gamma band (30-250 Hz). The neuromodulatory effect was also influenced by the prevalence of sharp wave ripples (SWR) during the pre-stimulation period, with firing rates modulated by up to 33% at units showing frequent baseline SWR.

**Conclusion:** Our findings suggest that AM and CW tFUS produce qualitatively different neuronal outcomes, and that baseline rhythms may effectively “gate” the response to tFUS.

## Introduction

Historically employed to image soft tissue, low-intensity focused ultrasound has more recently been shown to modulate brain activity (1; 2; 3; 4) in models spanning cell cultures (5; 6; 7), rodents (8; 9; 10; 11; 12; 13; 14; 15), primates (16; 17; 18; 19; 20), and humans (21; 22; 23; 24; 25; 26; 27; 28; 29; 30). Ultrasound overcomes the critical limitations of conventional (electromagnetic) non-invasive brain stimulation: it can be focused through the skull with millimeter precision (31) and penetrate deep brain regions (21). This raises the tantalizing possibility of utilizing transcranial focused ultrasound stimulation (tFUS) to modulate mesoscale neural circuits without the need for surgery, potentially providing novel interventions for the host of neurological and psychiatric disorders associated with aberrant brain activity (32).

To date, two general paradigms have been proposed for ultrasonic neuromodulation: continuous-wave (CW) and pulsed-wave (PW) tFUS. CW tFUS consists of a constant amplitude sinusoid, while PW tFUS is delivered as a series of tone bursts with a fixed duty cycle (33). In both cases, the sonication duration is generally short, most often in the tens to hundreds of milliseconds (2), although longer sonications have been found to yield long-lasting effects (16; 34; 10). There is not yet a clear consensus as to the relative efficacy of CW versus PW tFUS, with some studies finding that PW is able to elicit neuromodulation at lower intensities than CW (35), while others have reported a greater efficacy with CW tFUS (36). Given that the specificity of tFUS may originate from tailoring the ultrasonic dose (37) (i.e., frequency, intensity, duration, waveform), the identification of novel paradigms for ultrasonic neuromodulation is important to the advancement of tFUS as an intervention for disorders of the central nervous system.

Substantial variability in neural and behavioral outcomes has been widely reported in non-invasive brain stimulation (38; 39; 40), including focused ultrasound (28). Identifying the sources of this variability, whether it be exogenous (i.e., positioning of the transducer, anatomical differences) or endogenous (i.e., baseline neurophysiology) is essential to achieving robust and predictable outcomes with tFUS. The dynamics of the sonicated region leading up to stimulation, especially neural oscillations, may exert a causal influence on the subsequent response to stimulation (41). Electrophysiological brain rhythms may be readily captured with the electroencephalogram (EEG) or local field potentials (LFP). To our knowledge, however, the influence of baseline brain state on neuronal response to ultrasonic neuromodulation has not yet been investigated.

Here we propose and test a new mode of tFUS employing amplitude modulation (AM). Originating from telecommunications, AM is a signaling approach where the amplitude of a fast “carrier” wave is sinusoidally modulated at a relatively slow rate. In the context of tFUS, AM allows the embedding of a low frequency (analogous to the pulse repetition frequency in PW tFUS) into the ultrasonic stimulus, but in a smooth manner that avoids abrupt pressure transitions that may yield undesired effects during stimulation (15). To evaluate the effect of the proposed AM tFUS on neuronal activity, we stimulated the rat hippocampus while simultaneously recording electrophysiological responses from the sonicated region. We conditioned the resulting changes in spiking on baseline population activity, considering both the power of oscillations in canonical frequency bands as well as stereotyped markers of hippocampal excitability, namely sharp wave ripples (SWR) (42).

We found opposing effects on neuronal spiking with AM versus CW tFUS: multi-unit activity (MUA) was reduced with AM stimulation, while increasing with CW tFUS. Importantly, we also found a significant relationship between the power of endogenous brain rhythms prior to stimulation and the subsequent change in spiking. For both AM and CW stimulation, the firing rate change produced by tFUS was significantly larger when the pre-stimulation power of LFP oscillations was high, particularly in the gamma (30-250 Hz) band. Similarly, units with a high prevalence of baseline SWR responded more strongly to tFUS. Our findings suggest that AM tFUS represents a distinct paradigm for tFUS that may be well-suited to applications requiring the reduction of activity. More generally, the results underscore the importance of brain state in shaping the outcome of ultrasonic neuromodulation.

## Materials and Methods

Data were obtained from 18 adult male Long Evans rats weighing at least 350g (426.0 ± 32.2g, mean ± sd). All experimental procedures were approved by the Institutional Animal Care and Use Committee of the City College of New York, City University of New York.

### Transcranial focused ultrasound stimulation

In *n* = 8 rats, AM (40 Hz AM frequency, 100% AM depth, carrier frequency 2.0 MHz, sinusoidal) and CW waveforms (carrier frequency 2.0 MHz, sinusoidal) were generated with a waveform generator (Keysight 33500B Series). The output of the waveform generator was fed into the input of an RF Amplifier (Electronics and Innovation, 40W). The amplifier provided the drive voltage into a single-element ultrasonic transducer (Ultran KS25-2 immersion transducer, 2 MHz, 6.25 mm active diameter). Empirical measurements in a watertank indicated that the beampattern of this transducer has a full-width-half max (FWHM) of 2.4 mm laterally and 10 mm in depth. In the remaining *n* = 10 rats, AM tFUS (40 Hz AM frequency, 100% AM depth, carrier frequency 2.5MHz, sinusoidal) and CW tFUS (carrier frequency 2.5 MHz, sinusoidal) was delivered with an integrated FUS system (Sonic Concepts TPO-201) feeding a dual-channel, axial steered ultrasonic transducer (Sonic Concepts SU-132, 9.625 mm active diameter). The FWHM of this transducer was empirically measured as 1 mm lateral and 1.5 mm in depth (see Fig 1A). In all experiments, the transducer was mounted onto a micromanipulator arm of a stereotaxic frame (David Kopf Instruments) and coupled to the rat skull with ultrasonic coupling gel. The transducer was positioned over the desired anatomical target with the transducer face parallel to the skull (active stimulation: the right hippocampus −3.5 mm AP, +2.5 mm ML, targeted depth of 3.5 mm; sham stimulation: the left olfactory bulb +6 mm AP, +1.5 ML, targeted depth of 3.5 mm, coordinates relative to skull bregma and midline).

**Figure 1:**
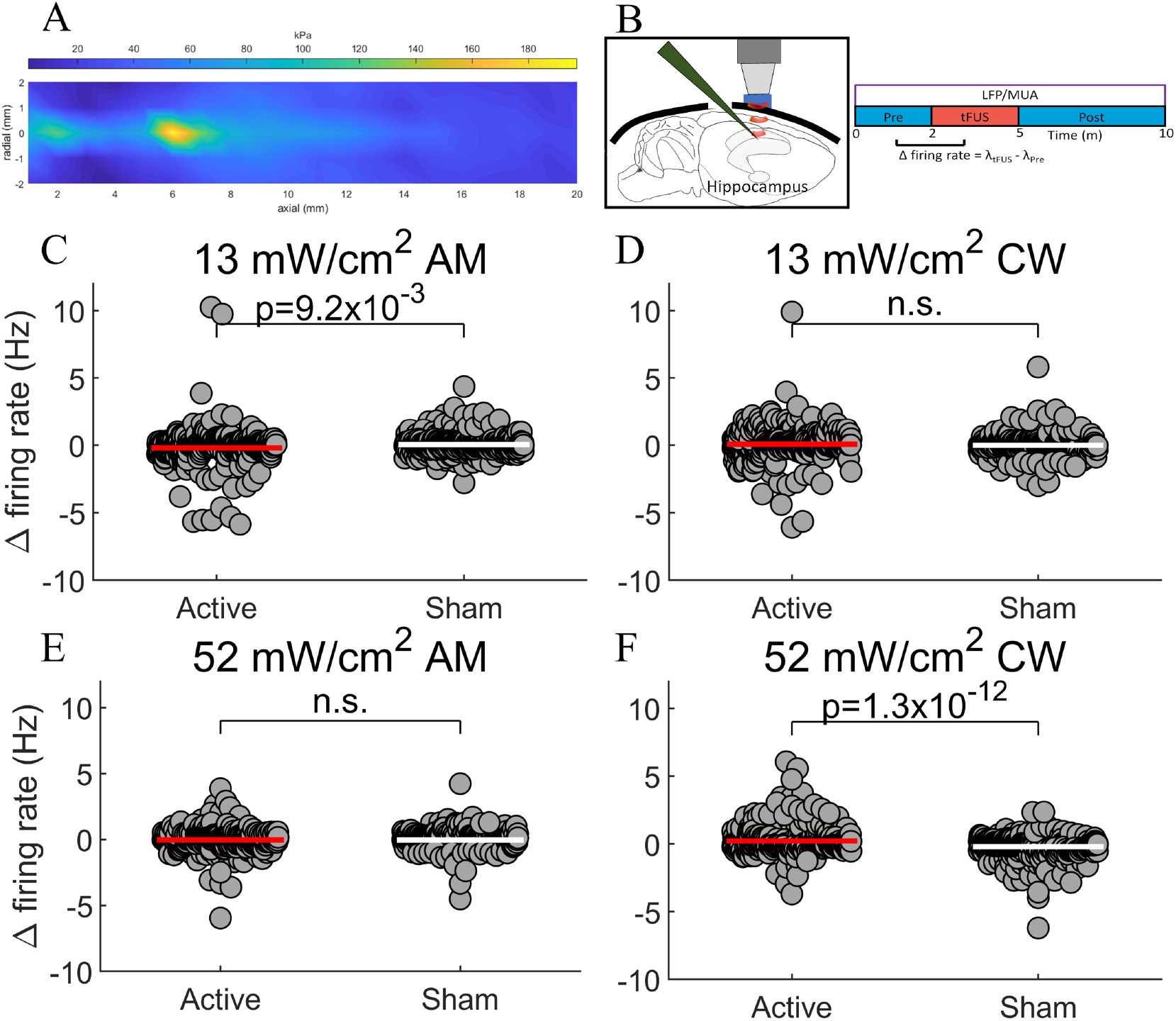
AM and CW tFUS produce opposing effects on firing rate. **(A)** The empirical beampattern of a transducer employed during the experiments, where the lateral and axial resolution are 1 and 1.5 mm, respectively. **(B)** Multi-unit activity (MUA) was captured from the hippocampus concurrently to AM and CW tFUS. We measured the change in firing rate observed during sonication relative to the preceding two minutes. **(C)** Compared to sham stimulation, 13 mW/cm^2^ AM tFUS significantly reduced firing rate (*p* = 0.0093, *n* = 392, t-test). No change in firing rate was resolved during stimulation with **(D)** 13 mW/cm^2^ CW or **(E)** 52 mW/cm^2^ AM tFUS. **(F)** On the other hand, a significant firing rate increase was found during 52 mW/cm^2^ CW tFUS (*p* = 1.3 × 10^-12^). The magnitude of both the reduction by AM and increase by CW was approximately 0.2 Hz, representing 10% of the baseline rate. These findings suggest that AM and CW tFUS yield distinct neuronal outcomes, and imply a complex interaction between ultrasonic intensity and waveform.

### Experimental design

A within-subjects design was employed where each animal received all eight experimental conditions, which spanned all combinations of two intensity values (13 mW/cm^2^ and 52 mW/ cm^2^), two waveforms (40 Hz AM and CW), and the two stimulation types (active and sham). The condition ordering was counterbalanced throughout the cohort to control for any order effects. A 50 minute interval was added between each condition to allow any outlasting effects to dissipate.

### Acoustic intensity calibration

Ultrasonic pressures were measured in a water tank with a calibrated hydrophone (Onda Corporation). Pressures were determined in both free field as well as with a model rat skull placed between the transducer and hydrophone. The values reported in the *Results* correspond to the intensities with the skull present. From the water tank calibrations, it was determined that the skull attenuates the acoustic pressure to a value that is 2/3 of the free-field pressure. For each transducer, the drive voltage required to produce acoustic intensities of 13 mW/ cm^2^ and 52 mW/ cm^2^ were determined and employed in the experiments. The pressure corresponding to 13 mW/ cm^2^ was 14 kPa, with an associated mechanical index (MI) of 0.01 at a 2 MHz center frequency. The pressure corresponding to 52 mW/ cm^2^ was 28 kPa, with an associated mechanical index of 0.02.

### Anesthesia and surgery

Prior to surgical experimentation, selected animals weighing at least 350g were fasted for 12-14 hours to increase urethane absorption. On the day of surgical experimentation, animals were placed into an induction chamber and induced with gaseous isoflurane at 3% (L/min). Animals were removed from the induction chamber and a nose cone was attached so that the dorsal hair could be shaved in preparation for the craniotomy. Animals were then placed on a stereotaxic frame (David Kopf Instruments) with earbars securing the head. The isoflurane concentration was then reduced to 2% (L/min). A cross incision was performed over the dorsal skull to expose the cranium and skull landmarks. The distance from bregma to the interaural line was measured so that AP coordinates could be adjusted to account for differences in animal size. The center of the forthcoming craniotomy was marked at −7.5 AP and +2.5 ML (directly posterior of the placement for active stimulation). This allowed sufficient clearance between the edge of the transducer and the border of the craniotomy. A 2 mm by 2 mm craniotomy was performed over the marked area, followed by the removal of the dura. A small titanium screw was implanted into the skull (left hemisphere) to provide an electrical ground for the electrophysiological recordings. The concentration of isoflurane was further reduced to 1% (L/min) and a urethane cocktail (1.5g/kg diluted with 2.5ml/g saline, divided into 3-4 doses with one dose administered every 10 minutes) was administered via intraperitoneal injection. After the final urethane injection, isoflurane was again lowered to 0.5% (L/min) to allow for urethane absorption and anesthesia transition. After 30 minutes, isoflurane was discontinued and a 120 minute period was allowed for complete expulsion of isoflurane and to achieve a stable anesthesia plane prior to the experiment.

### Electrophysiology

Multi-unit activity (MUA) and local field potentials (LFP) were recorded with a linear 32-channel silicon electrode array (NeuroNexus A32, 100 *μ*m spacing between adjacent contacts). Signals were recorded with a digital acquisition system (NeuroNexus SmartBox) at a sampling rate of 30 kHz. The probe was placed into the center of the craniotomy at an angle of 53° from vertical (angled towards the posterior), and then advanced 6 mm so that the contacts sampled multiple subregions of the hippocampal formation, including CA1, CA3, and the dentate gyrus. Electrophysiological recording commenced two minutes before the onset of ultrasonic stimulation, continued throughout the three-minute stimulation period as well as an additional five minutes post-stimulation, resulting in 10 minute data recordings for each experimental condition. A continuous trigger was outputted from either the waveform generator or integrated FUS system to the digital acquisition system throughout ultrasonic stimulation to mark tFUS onset.

### MUA analysis

We employed the Kilosort 2 (43) software running on Matlab (Mathworks, Release 2019a) to perform automated spike detection and sorting. This technique forms a generative model of the extracellular voltage, learns a spatiotemporal template of each spike waveform based on the singular value decomposition, and employs multiple passes through the data to yield clusters of spikes. We employed most default parameters provided by the developers, as described in the standard configuration file provided at github.com/MouseLand/Kilosort. We employed a high pass filter cutoff of 400 Hz to exclude slow activity, lowered the detection threshold from −6 to −5 standard deviations, and doubled the default batch size to improve the learning algorithm for the spatiotemporal template. Spike detection and sorting was performed separately for each animal, and all 8 experimental recordings were chronologically assembled into a single data record prior to processing. The results of the automated procedure were imported into the Phy software (44), an open source Python library and graphic user interface for manual curation of large-scale electrophysiological data. Visual inspection of each identified unit’s auto- and cross-correlograms, distribution of amplitudes, and spike waveforms was performed to discard units deemed to be non-biological (*n* = 196 units were discarded). The analysis led to a total of *n* = 392 units whose firing rates were then probed for changes due to the tFUS intervention.

### LFP analysis

LFP signals were analyzed offline using custom Matlab scripts (Matworks, Release 2019a). Data was bandpass filtered to the 1-250 Hz band with a second-order Butterworth filter and then downsampled to 500 Hz. A series of notch filters were then applied to remove 60 Hz noise and its first four harmonics. Robust principal components analysis (robust PCA) (45) was employed to remove gross artifacts by decomposing the observed data matrix into low-rank and sparse components. Due to the smoothness of volume conducted signals, the sparse component is expected to be artifactual and was thus removed from the data. Data were then transformed into the frequency domain by performing Thomson multi-taper spectral analysis with a time-bandwidth product of 4. All spectra were normalized by the total spectral power measured in the pre-stimulation period of each animal’s first condition to account for varying power levels between animals. Spectra were further resampled in the frequency domain to 1000 samples between 1 and 250 Hz via linear interpolation. Spectral powers for any frequency greater than 57 and less than 63 hz were marked as missing data due to possible contamination from line noise. Spectral powers were then logarithmically transformed, averaged across contacts within the hippocampus (two contacts outside of the hippocampus were excluded), and then averaged across the appropriate segment of time (baseline, stimulation, post-stimulation). The dependent variable was formed as the difference in LFP power between the stimulation (or post-stimulation) and baseline segments, measured frequency-wise.

### Baseline LFP measurement

In order to investigate the role of baseline brain state on neuromodulation outcome, we computed the the mean power in the following frequency regions: delta (1-3 Hz), low theta (4-6 Hz), high theta (6-10 Hz), and gamma (30-250 Hz). For each MUA unit, we determined the electrode contact best expressing the spikes by searching for the channel with largest spike amplitude. We then measured the logarithmically transformed LFP power during the two minute pre-stimulation period at the identified contact. This formed the independent variable whose influence on firing rate changes were probed throughout the main text. When delineating “low” and “high” baseline LFP power, we partitioned the units into two groups with assignment based on the LFP power relative to the median power across all units. This assignment was performed separately for each frequency band.

### Sharp wave ripple detection

In order to measure the prevalence of sharp wave ripples (SWRs) during the pre-stimulation period, we followed the detection algorithm described previously by Levenstein et al (46). Sharp waves were detected when the power of the contact-averaged LFP in the 2-50 Hz band exceeded 2.5 standard deviations of the mean power. Only segments exceeding 20 ms were retained. Ripples were detected when the contact-averaged LFP power in the 80-249 Hz band exceeded 2.5 standard deviations of the mean power, with a minimum segment length of 25 ms. Samples that passed both the sharp wave and ripple detectors were then designated as SWRs. The proportion of time “spent” in SWR then served as the independent variable in the subsequent analysis of the gating of firing rate by the prevalence of SWR. In order to define periods of rare and frequent SWR, we performed a median split on the percentage of samples passing the SWR detector. Units that did not show any SWRs were excluded from the analysis. This resulted in a sample size of *n* = 189 to *n* = 270 units, depending on the condition.

### Statistical Testing

Unless otherwise specified, testing for statistical significance was carried out by comparing the dependent variable measured under active stimulation against that observed with sham stimulation. When testing for significant differences in firing rates, we employed paired t-tests. Where appropriate, correction for multiple comparisons was conducted by controlling the false discovery rate (FDR) at 0.05. When probing significant discrimination of firing rate change by the baseline LFP power (Fig 4), we performed a non-parametric test scrambling the assignment of non-responders (firing rate change less than the median) and responders (firing rate change greater than the median). A total of 1000 mock data records modeling the null distribution of AUROC were then employed to test for significance of the true AUROC at each frequency (corrected for 2049 comparisons using the FDR). When probing tFUS-induced changes to the LFP power spectrum during and after stimulation, a non-parametric test scrambling the assignment of active and sham conditions was employed, with 1000 permutations employed to estimate the null distribution of LFP power spectral changes at each frequency. Cluster correction (47) was then employed to correct for multiple comparisons.

## Results

In order to determine the effects of AM tFUS on hippocampal neuron spiking, as well as its relationship to baseline neural activity, we recorded multi-unit activity (MUA) and local field potentials (LFP) from the rat hippocampus before, during, and after the application of 180 s of tFUS at two different intensities (*I*_spta_=13 mW/ cm^2^, *I*_spta_=52 mW/ cm^2^) and two waveforms (40 Hz AM, CW). The empirical beampattern of a transducer employed during the experiments is depicted in Fig 1A, where the lateral and axial full-width-half max (FWHM) are 1 and 1.5 mm, respectively. Each of the four doses was paired with a corresponding sham condition during which we stimulated the contralateral olfactory bulb to account for non-specific effects such as spontaneous activity changes and potential bone conduction of the ultrasonic stimulus. The order of all eight conditions was counterbalanced across the *N* = 18 animals tested. Semi-automated spike sorting identified a total of *n* = 392 MUA units. The primary outcome measure was the change in firing rate observed during tFUS relative to the baseline period (Fig 1B). We then related the observed changes in spiking rate to several LFP-derived markers of brain state, such as the power of oscillations and prevalence of sharp-wave ripples (SWRs) during the pre-stimulation period.

### AM and CW produce opposing effects on firing rate

We found a significant reduction in firing rate during 13 mW AM tFUS (Fig 1C: active firing rate change = −0.22 ± 0.087 Hz, sham firing rate change = 0.026 ± 0.031 Hz, means ± sem, *n* = 392 units from *N* = 18 animals, paired t-test, *p* = 0.0093). We also found a significant increase in firing rate during 52 mW CW tFUS (Fig 1F: active firing rate change = 0.21 ± 0.046 Hz, sham firing rate change= −0.21 ± 0.037 Hz, *p* = 1.38 × 10^-12^). We were not able to resolve a significant change in firing rate during either 13 mW CW tFUS (Fig 1D, *p* = 0.27) or 52 mW AM tFUS (Fig 1E, *p* = 0.94). To quantify the magnitude of the average firing rate changes observed during tFUS (approximately 0.2 Hz), we note that the mean baseline firing rate across all animals and conditions was 2.32 Hz, meaning that the effect of tFUS for both AM and CW waveforms was in the order of 10%.

In order to probe outlasting firing rate changes, we compared the mean firing rate observed prior to tFUS with that measured in the five-minute period immediately following stimulation. We found a marginal but significant effect of reduced firing rate for 13 mW AM tFUS (data not shown; active firing rate change = −0.17 ± 0.036 Hz, sham firing rate change = 0.087 ± 0.048 Hz, *p* = 0.040). There were no significant differences in firing rate during the five-minute period following tFUS for 13 mW CW, 52 mW AM, and 52 mW CW tFUS (Fig S1B-D, all *p* > 0.18).

To gain insight into the dynamics of the firing rate changes detected during tFUS, we measured spiking rate in a time-resolved manner, employing non-overlapping 15 second increments (Fig 2). We focused the analysis on the 13 mW AM and 52 mW CW doses, as these exhibited a significant change during stimulation when aggregating across time segments (Fig 1). For each 15 second window, we measured firing rate and tested for significant deviations from the baseline rate (stimulation onset occurred at 120 s). Compared to sham stimulation (Fig 2B), 13 mW AM tFUS produced extensive periods of significantly reduced firing rate during and after stimulation, extending 210 seconds into the post-sonication period (Fig 2C: significance during windows beginning at 120, 165-225, 315-360, 390, 435 and 495 s; during tFUS: *p* < 0.016, after tFUS: *p* < 0.013, paired t-tests of the difference between time-resolved firing rate and the baseline firing rate, *n* = 392, corrected for multiple comparisons by controlling the false discovery rate at 0.05). On the other hand, the increased firing rate observed during 52 mW CW tFUS was mostly confined to the stimulation period (Fig 2F: significance during all windows between 120 and 315 s; during tFUS: *p* < 1.8 × 10^-4^, after tFUS: *p* < 0.0055). Interestingly, for both 13 mW AM and 52 mW CW, the peak acute effect occurred during the 180-195 s window (i.e., at the onset of the second minute of sonication), implying a slow accumulating effect. To that end, we probed short-term time-locked changes at tFUS onset and did not observe any significant effects (data not shown).

**Figure 2:**
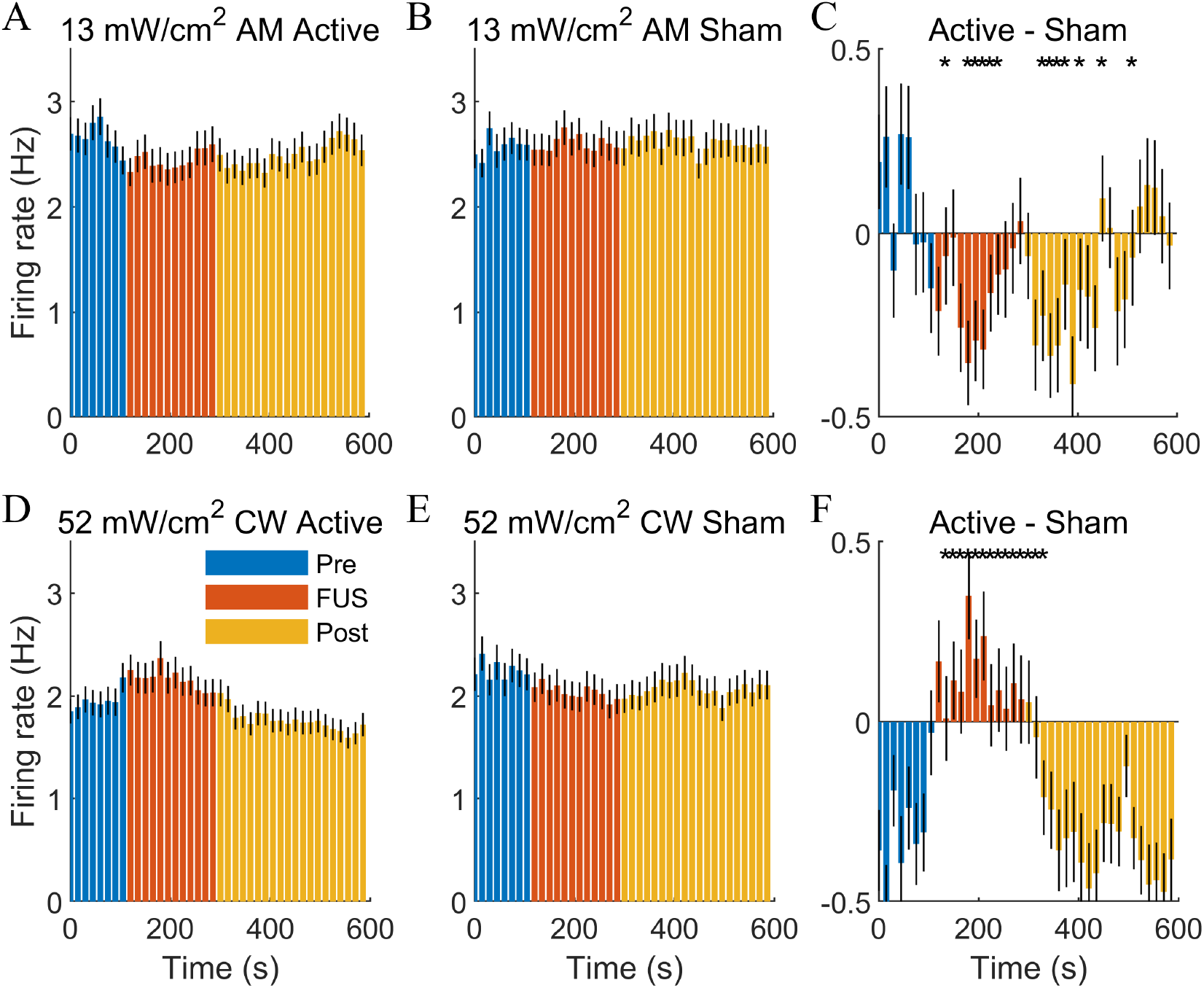
The time course of firing rate modulation suggests an accumulating effect. **(A)** Vertical axis depicts the mean firing rate in non-overlapping 15-second windows spanning the 10-minute recording period of the 13 mW/cm^2^ AM tFUS condition (baseline: green, stimulation: red, post-stimulation: orange). A reduction during and immediately after sonication is apparent in several time windows. **(B)** Time course of firing rate for sham AM tFUS. **(C)** The difference in firing rate between active and sham stimulation shows that the maximal reduction occurs during the second minute of sonication (i.e., 180 - 195 s into the recording), and is sustained for more than three minutes after sonication (i.e., 510 s). Asterisks denote a significant change relative to sham for the time window (during tFUS: *p* < 0.016, after tFUS: *p* < 0.013, t-test, *n* = 392, corrected for multiple comparisons by controlling the false discovery rate at 0.05). **(D)** Same as **(A)** but now for 52 mW/cm^2^ CW tFUS. Firing rate is increased during the stimulation period. **(E)** Same as D but now for sham stimulation. **(F)** The difference in firing rate between active and sham stimulation, where the peak effect is again observed 180-195 s into the recording. Unlike AM tFUS, significant changes in firing rate were mostly confined to the sonication period (during tFUS: *p* < 1.8*x* 10^-4^, after tFUS: *p* < 0.0055).

### tFUS neuromodulation is gated by baseline LFP power

We suspected that the effect of tFUS on spiking is influenced by the state of the neuronal population prior to stimulation, in particular the level of synaptic activity. To test this hypothesis, we measured the LFP power spectrum during the two-minute baseline period leading up to stimulation for each unit, calculating the strength of neural oscillations at the contact that most strongly registered the spike (see *Methods*). We then sought to determine whether the units exhibiting high levels of pre-stimulation oscillations would respond more strongly to tFUS. To that end, we partitioned units into two groups, those exhibiting relatively low LFP power (less than median) and those exhibiting high power levels (greater than median). We performed this classification separately for the delta (1-3 Hz), low theta (4-6 Hz), high theta (6-10 Hz), and gamma (30-250 Hz) frequency bands. If the LFP power modulates or “gates” the response to tFUS, we would expect the conditional distribution of firing rate to vary significantly between the two groups.

When considering all units, 13 mW AM tFUS reduced firing rate by an average of 0.22 Hz (Fig 1C). These reductions were significantly amplified at units showing elevated power in all four frequency bands. For units exhibiting low delta power, the firing rate change was only −0.040 ± 0.11 Hz; on the other hand, units whose delta power was above the median responded more prominently, with a firing rate change of −0.39 ± 0.14 Hz (Fig 3A, *p* = 0.043, *n* = 196, unpaired t-test). Moreover, the trend of stronger firing rate changes held for 3-6 Hz theta (Fig 3C; low: 0.017 ± 0.033 Hz, high: −0.45 ± 0.17 Hz, *p* = 0.0077), 6-10 Hz theta (Fig 3E: low: −0.0010 ± 0.034 Hz, high: −0.43 ± 0.17 Hz, *p* = 0.014), and gamma power (Fig 3G; low: −0.025 ± 0.047 Hz, high: −0.41 ± 0.17 Hz, *p* = 0.029). Note that, in all cases, the overall effect of 13 mW AM tFUS on firing rate was contributed by the units with high pre-stimulus power (i.e., an all-or-nothing phenomenon emerged, in that units with low oscillatory power did not experience any change).

**Figure 3:**
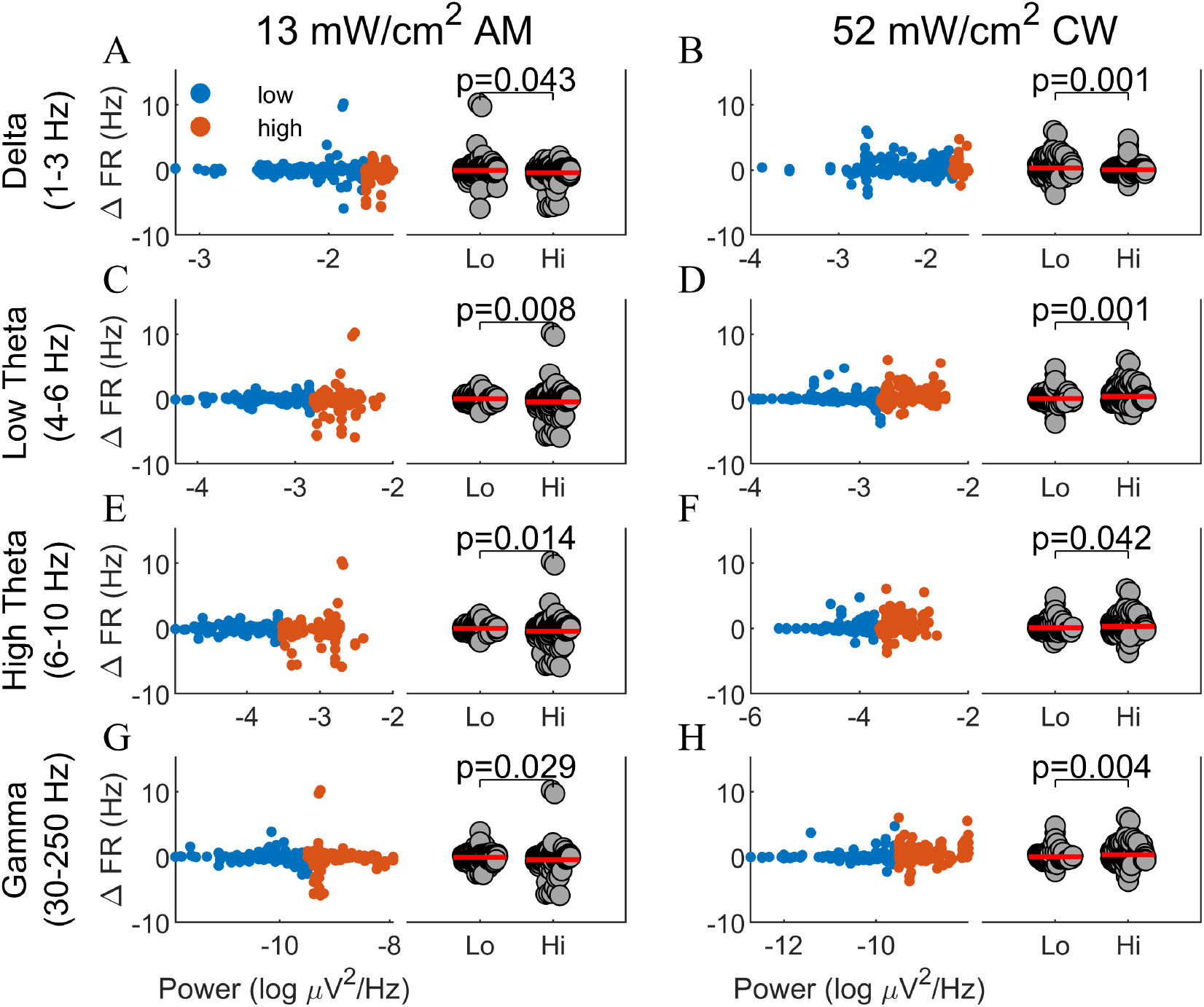
Firing rate changes are gated by pre-stimulation LFP power. In order to test the hypothesis that the brain state leading up to sonication influences the subsequent response to tFUS, we measured baseline LFP power in the delta (1-3 Hz), low theta (4-6 Hz), high theta (6-10 Hz), and gamma (30-250 Hz) frequency bands. For each frequency band, units were then partitioned into two groups: those whose baseline power was less than the median (blue) and those whose power fell above the median (red). **(A)** The firing rate induced by AM tFUS was significantly stronger when preceded by relatively high delta power (*p* = 0.043, *n* = 196, t-test, corrected for 8 comparisons by controlling the FDR at 0.05). **(B)** Conversely, the increase in firing rate due to CW tFUS was significantly larger when preceded by *low* delta power (*p* = 0.001). **(C-D)** For both AM and CW, the magnitude of the firing rate change was significantly larger when the period leading up to sonication was marked by higher power in the 4-6 Hz theta band (AM: *p* = 0.008, CW: *p* = 0.001). **(E-F)** Same as (C-D) but now for the 6-10 Hz portion of the theta band. The firing rate reductions and increases from AM and CW tFUS, respectively, were larger when preceded by higher baseline 6-10 Hz power (AM: *p* = 0.014, CW: *p* = 0.042). **(G-H)**. The power of gamma band (30-250 Hz) oscillations similarly predicted the size of the firing rate change for both AM (*p* = 0.029) and CW tFUS (*p* = 0.004). Thus, in all conditions, the strength of pre-stimulation rhythms was predictive of the subsequent change in spiking from tFUS.

A very similar finding was observed for 52 mW CW tFUS. In this condition, however, *low* delta power was associated with a more prominent response to tFUS: 0.35 ± 0.079 Hz when the delta power was below the median, and only 0.063 ± 0.043 Hz when the delta power was above the median (Fig 3B: *p* = 0.0013). As was the case with 13 mW AM tFUS, high levels of 4-6 Hz theta (Fig 3D; low: 0.053 ± 0.050 Hz, high: 0.37 ± 0.075 Hz, *p* = 0.0005), 6-10 Hz theta (Fig 3F; low: 0.12 ± 0.047 Hz, high: 0.30 ± 0.078 Hz, *p* = 0.042), and gamma (Fig 3H; low: 0.077 ± 0.044 Hz, high: 0.34 ± 0.079 Hz, *p* = 0.0037) powers were associated with a significantly stronger response to tFUS.

In order to gain greater insight into the relationship between pre-stimulation rhythms and the subsequent response to tFUS, we visualized the empirical joint distribution between pre-stimulus LFP power (horizontal axis) and firing rate change (vertical axis) due to tFUS (left panels in Fig 3). It became evident that units at the low end of the pre-stimulation power often exhibited no response. Moreover, the likelihood of a response generally increased as the baseline LFP power increased, with the firing rate change exhibiting a greater variance with increasing baseline LFP power. Note that this increased variance extended in both directions, in that some units exhibited a marked change in firing against the general trend (e.g. Fig 3E-F). These findings suggest that increased synaptic activity, as reflected in greater LFP power, signifies a higher general sensitivity to mechanical stimulation.

To identify the frequencies that are most predictive of successful neuromodulation, we considered the prediction of firing rate change from the LFP power at each individual frequency. Given two distributions (i.e., LFP power for units with a strong response, LFP power for units with a weak response), the Receiver Operating Characteristic (ROC) curve provides a cumulative measure of the spread between the two distributions. For this analysis, we binarized the firing rate change by comparing each unit’s change to the median, effectively partitioning the units into “responders” and “non-responders”. Integrating the ROC curve yields an aggregated statistic termed the Area under the ROC curve (AUROC). Values near 0.5 signify no discrimination between distributions, while values closer to 0 or 1 denote that the predictor (i.e., LFP power) has a strong influence on the target (i.e., binarized firing rate change). For 13 mW AM tFUS, a significantly negative AUROC was observed in punctate regions of the delta and theta bands, as well as an extensive portion of the gamma frequencies (Fig 4A: significant frequencies indicated with markers above the horizontal axis, permutation test, corrected for multiple comparisons by controlling the false discovery rate at 0.05). Given that AM tFUS decreased firing rate, a value of AUROC less than 0.5 signifies that units with high power in these bands responded with a more negative firing rate change. For 52 mW CW tFUS, a region of significant discrimination was found at very low frequencies (Fig 4B, trough near 2 Hz), in addition to a broad region of significance spanning frequencies from 3 to 140 Hz (Fig 4B). This indicates that in CW tFUS, high LFP power in the theta and gamma region, as well as low delta power, promotes successful modulation of firing rate.

**Figure 4:**
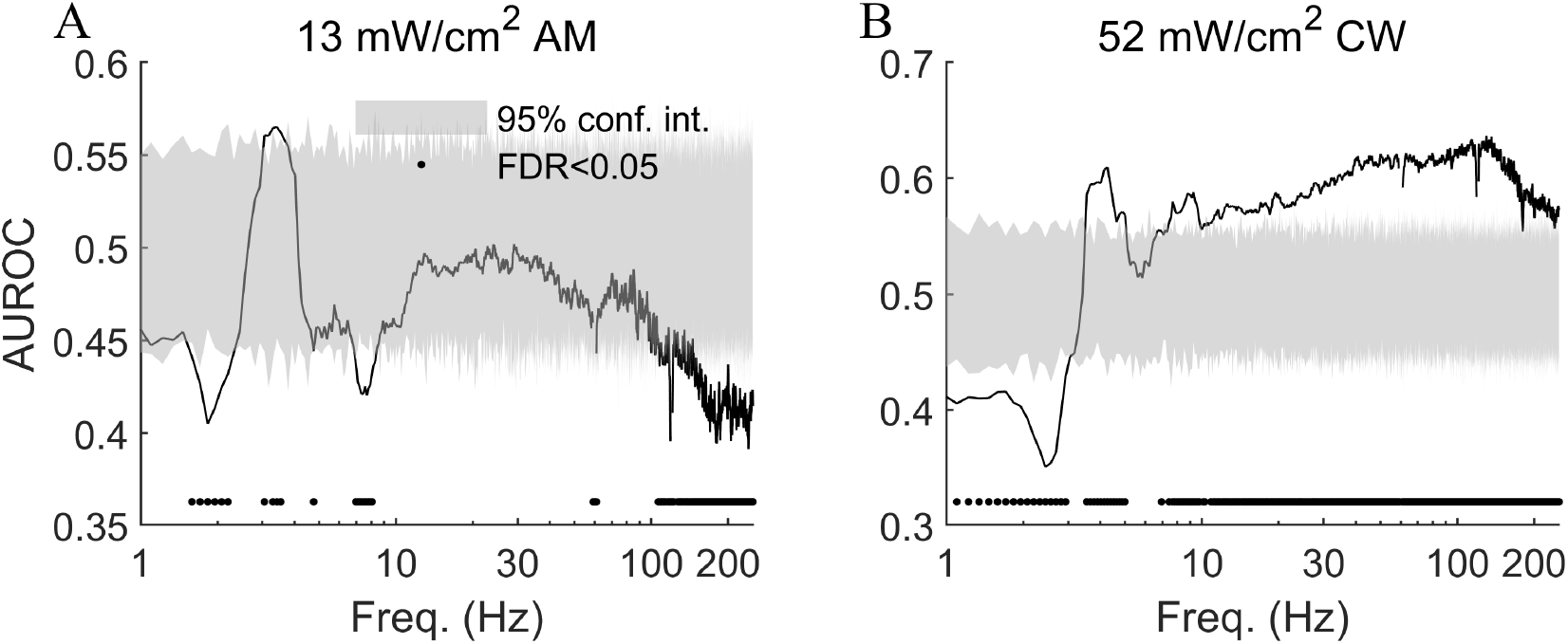
The power of pre-stimulation gamma oscillations is predictive of neuronal response to tFUS. In order to identify the LFP frequencies most predictive of neuromodulation outcome, we performed linear discriminant analysis aimed at classifying “responders” (units showing a firing rate change greater than the median) from “non-responders” (units whose firing rate changed less than the median) from the baseline LFP power at a given frequency (horizontal axes). The Area under the Receiver Operating Characteristic curve (AUROC, vertical axes) is a measure of the separation between distributions of baseline LFP power for responders versus non-responders. **(A)** For AM tFUS, we found significant discrimination in punctate regions of the delta and theta regions, and an extensive region of significant discrimination in the 100 - 250 Hz gamma region. The shaded grey region indicates the 95% confidence interval for AUROC under the null hypothesis. Due to AM stimulation reducing firing rate, AUROC values less than 0.5 denote that high values of gamma power led to more negative changes (i.e., a stronger neuromodulation). **(B)** For CW tFUS, low values of delta power (1 - 3 Hz) and high values of both theta and especially gamma (30 - 250 Hz) power led to a greater increase in firing rate during tFUS. These findings indicate that baseline gamma band activity is a strong predictor of neuronal sensitivity to ultrasonic neuromodulation.

### tFUS neuromodulation is enhanced during periods of SWR

Under conditions of urethane anesthesia and natural sleep, the hippocampus alternates between periods of spiking and inactivity. The active periods manifest in the LFP by sharp wave ripples (SWRs), short bursts of high-frequency activity (42; 46). Given that SWRs denote an excitable state, we suspected that tFUS would result in stronger responses when applied during periods of frequent SWR. To test this, we separated units into two categories: those whose baseline periods exhibited fewer than the median number of observed SWRs, and those that showed more than the median.

When comparing the outcome of tFUS between the two groups, we found large effects for all tFUS conditions. With 13 mW AM tFUS, the firing rate reduction at units marked by frequent SWR was very pronounced: −1.11 ± 0.32 Hz, significantly larger than the change at units with rare baseline SWR: 0.11 ± 0.05 Hz (Fig 5A; *p* = 2.58 × 10^-5^, *n*_low_ = 169, *n*_high_ = 101, unpaired t-test). The mean basal firing rate for units with prevalent SWR was 3.34 ± 0.40 Hz, meaning that the spiking rate reduction for these units was approximately 33%. Interestingly, the influence of baseline SWR was also significant for 13 mW CW tFUS, with the frequent SWR state leading to a firing rate reduction of −0.28 ± 0.10 Hz and rare SWR leading to an increase of 0.51 ± 0.12 Hz (Fig 5B: *p* = 1.52 × 10^-6^). For 52 mW AM tFUS, frequent baseline SWR led to a firing rate reduction of −0.35 ± 0.079 Hz, while units with few SWRs did not experience a modulation: 0.021 ± 0.030 Hz (Fig 5C; *p* = 2.23 × 10^-6^). Finally, for 52 mW CW tFUS, stimulating during periods of frequent SWR led to a significant increase in firing rate relative to rare SWR (Fig 5D: frequent: 0.54 ± 0.099 Hz, rare: 0.074 ± 0.049 Hz, *p* = 4.12 × 10^-6^). The sensitivity of the tFUS effect to the prevalence of SWRs suggests that low-intensity ultrasound’s modulation interacts strongly with concurrent synaptic input into the sonicated region.

**Figure 5:**
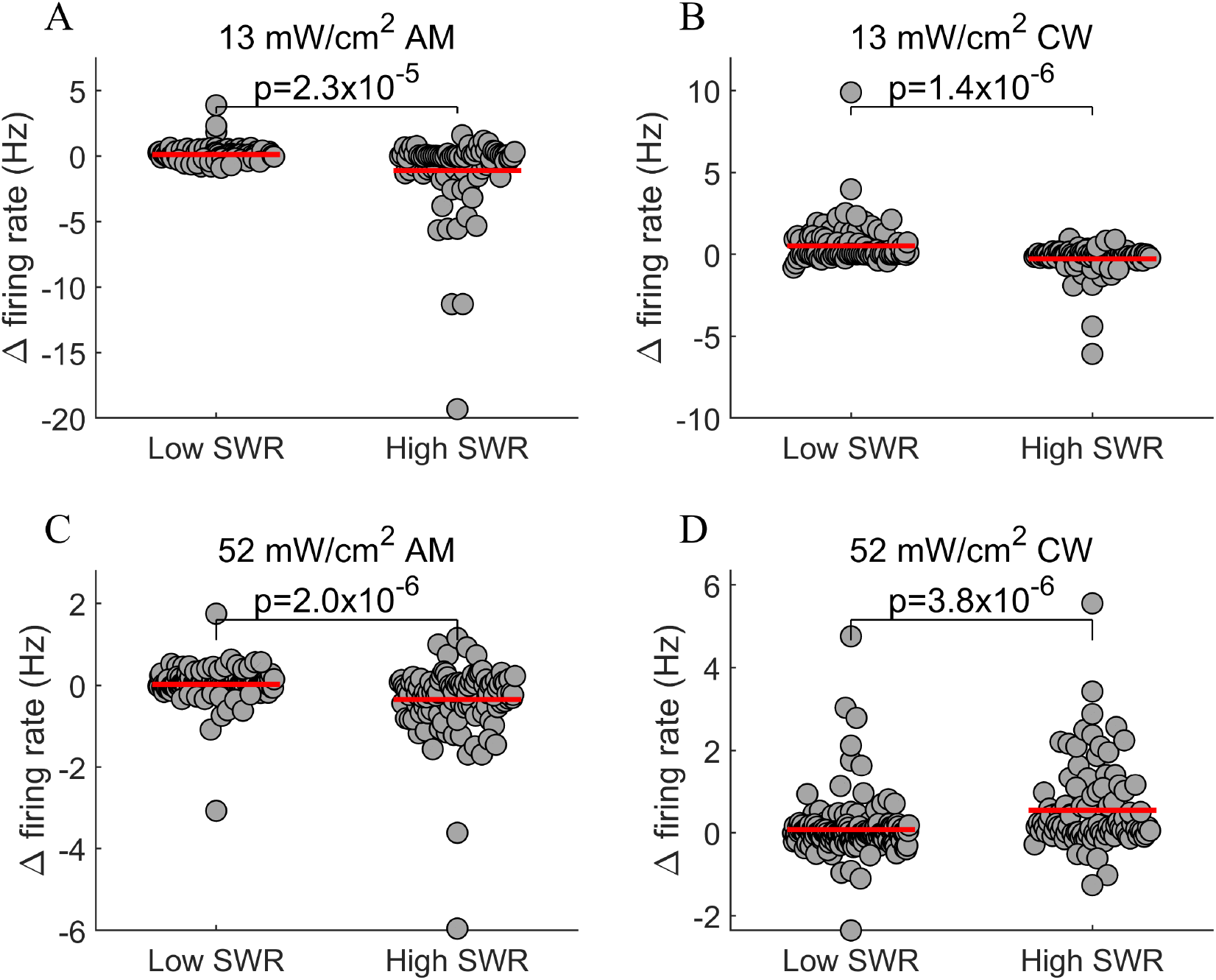
The prevalence of SWR in the pre-stimulation period is a strong driver of neuromodulation outcome. Sharp-wave ripples (SWR) in the hippocampal LFP reflect a transient state of increased neural excitability. To test whether neuronal sensitivity to tFUS is modulated by the prevalence of SWR leading up to sonication, units were partitioned into two groups depending on the amount of SWR during the baseline period (i.e., median split). **(A)** Stimulation with 13 mW AM tFUS produced a significantly larger reduction in firing rate (i.e., 33%) when applied during frequent SWR (*p* = 2.58 × 10^-5^, *n*_low_ = 169, *n*_high_ = 101, t-test). **(B)** The prevalence of baseline SWR also modulated the change in firing rate observed with 13 mW CW tFUS (*p* = 1.52 × 10^-6^). **(C)** Firing rate reductions during 52 mW AM tFUS were significantly larger in units with frequent SWR (*p* = 2.23 × 10^-6^). **(D)** Units with more frequent SWR also exhibited a larger increase in firing due to 52 mW CW tFUS (*p* = 4.12 × 10^-6^). These findings suggest that the success of tFUS is linked to the presence of concurrent synaptic input at the sonicated region.

### tFUS modulates theta and gamma power

The LFP provides a complementary measure of neural activity, reflecting the overall synaptic activity at the recorded region. We probed changes in LFP power during and after sonication. Consistent with the findings of the MUA analysis, significant changes were resolved in the 13 mW AM tFUS and 52 mW CW tFUS conditions. In particular, a significant increase of gamma power was observed during 13 mW AM tFUS (Fig 6A, significant cluster from 63-94 Hz, *p* < 0.05, permutation test, cluster corrected for multiple comparisons). An increase in both theta and gamma bands was observed during and also after 52 mW CW tFUS (Fig 6D, significant clusters at 8-18, 81-90, 105-145, 159-193, 203-215, 224-229 Hz). No significant LFP power changes were resolved at 13 mW CW or 52 mW AM tFUS (Fig 6B-C).

**Figure 6:**
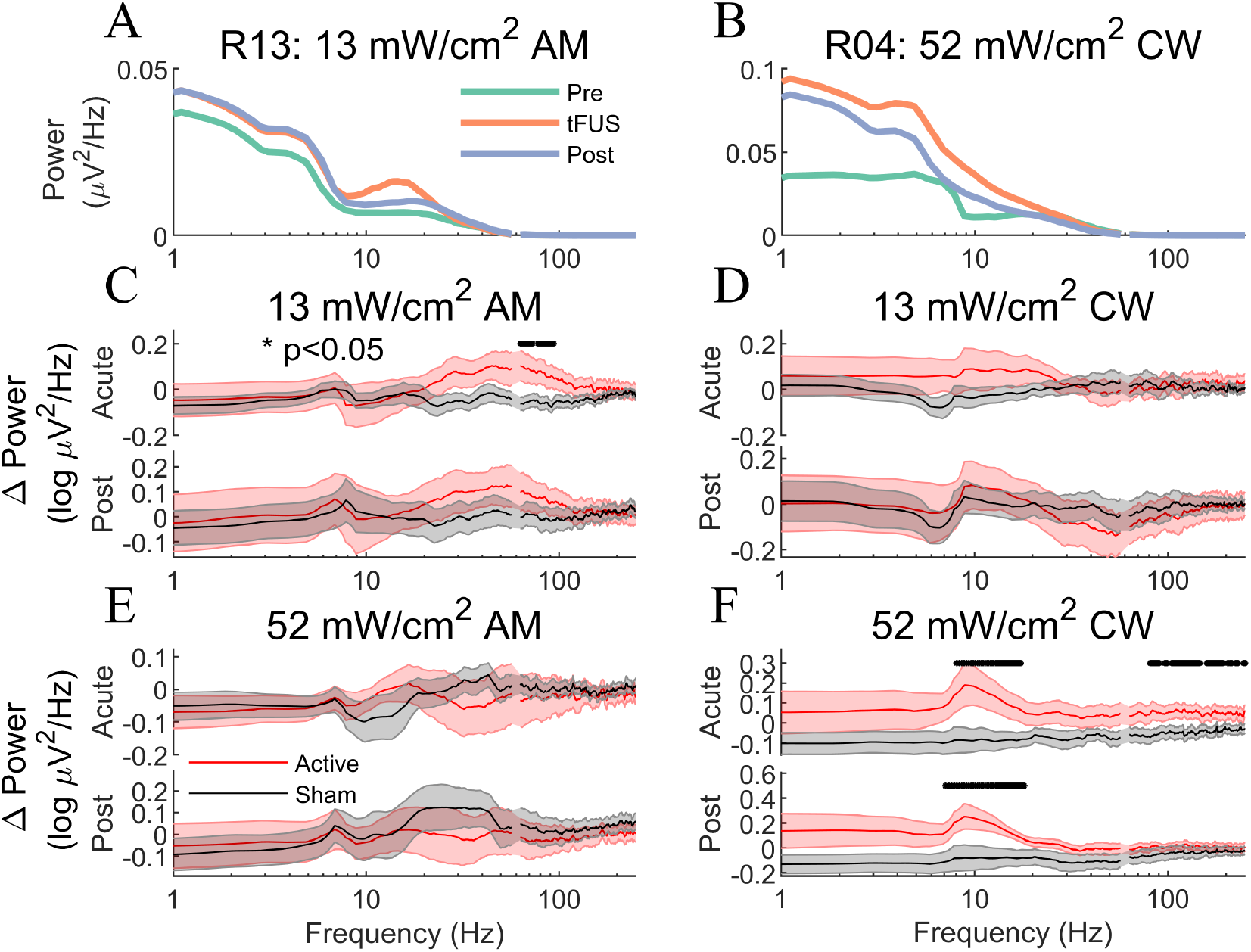
tFUS modulates LFP power in the theta and gamma bands. To determine whether tFUS modulates overall levels of synaptic activity, we probed changes in LFP power during and after sonication. **(A)** LFP power spectrum before, during, and after 13 mW/cm^2^ AM tFUS for a sample animal. **(B)** Same as **(A)** but now for a sample animal at 52 mW/cm^2^ CW tFUS. **(C)** At the group level, 13 mW/cm^2^ AM tFUS significantly increased power in a cluster of gamma band frequencies (63-94 Hz, *p* < 0.05, *n* = 18, permutation test, cluster corrected for multiple comparisons). The increased gamma power was not sustained following sonication. Mirroring the MUA findings, no significant LFP power changes were detected for either **(D)** 13 mW/cm^2^ CW or **(E)** 52 mW/cm^2^ AM tFUS. **(F)** On the other hand, a significant increase in both theta (8-18 Hz) and high gamma (80-229 Hz) was identified during 52 mW/cm^2^ CW tFUS (*p* < 0.05), with the theta band increase persisting beyond the stimulation period.

## Discussion

Neuromodulation techniques capable of directly evoking neuronal firing are referred to as “super-threshold”. Examples of these are transcranial magnetic stimulation, deep brain stimulation, and optogenetics. On the other hand, sub-threshold techniques such as transcranial direct current stimulation (tDCS) do not directly evoke firing but rather bring the membrane potential closer to (or further from) the threshold for action potential initiation. Our findings suggest that, at the intensities tested here (13 - 52 mW/ cm^2^), tFUS belongs to the subthreshold category. We found no evidence of time-locked firing, and the largest changes in firing rate were observed during the second minute of a three-minute sonication, implying an accumulating effect. Moreover, the large influence of pre-stimulation LFP oscillations and SWRs on the resulting neuromodulation outcome suggest that concurrent synaptic input is a key ingredient of successful neuromodulation. LFP power reflects the amount of coherent synaptic input into the region (48), while SWRs are highly synchronous events marked by coordinated firing across many neurons (42). Notably, SWRs are associated with a transient increase in hippocampal excitability (49), consistent with our finding of an enhanced response to tFUS during periods of frequent SWR. Both AM tFUS, which reduced firing, and CW tFUS, which increased it, led to significantly larger effects on spiking in the presence of strong LFP rhythms and frequent SWRs. The term “gating” denotes that without a sufficient level of synaptic drive into the stimulated cell, tFUS may not produce a change in spiking rate. Motivated by similar hypotheses in electrical stimulation, one approach that has been employed in tDCS is to pair the stimulation with a task that engages the stimulated area (50). Future tFUS studies that combine behavioral interventions with stimulation may further clarify the role of concurrent input on neural outcomes. Note that in contrast to what was found here, earlier investigations of tFUS reported very short latency responses (8) that are more consistent with a super-threshold mechanism, and these have been attributed by some as being influenced by an auditory confound (12; 13). Furthermore, it is possible that at higher acoustic intensities, the effect of tFUS may become super-threshold, and in particular in the event that a local temperature increase is produced at the sonicated region.

The majority of recent tFUS investigations have employed brief sonications, typically in the tens to hundreds of milliseconds (2), with stimulation applied between relatively long intertrial intervals. An advantage of this approach is that it affords an increase in statistical power, as the evoked response may be time locked and averaged over many repeated trials. Nevertheless, previous investigations that have instead employed single sonications with long duration have reported outlasting effects. For example, 40 seconds of tFUS to the primate brain was found to produce a long-lasting effect on functional connectivity (16). A 20-minute application of tFUS in the rat was shown to shorten the time required to recover from ketamine-xylazine anesthesia (51). Three minutes of tFUS was shown to reduce epileptic discharges when applied after the onset of chemically-induced seizures (34). Similarly, here we applied 180 s of tFUS, and indeed were able to find a significant modulation beyond the sonication period. The outlasting effects were more prominent with AM tFUS, where firing rate was significantly reduced as far as four minutes into the post-sonication period. In this way, our finding supports the notion that relatively long sonication periods promote sustained neuromodulation, which will be required for future clinical investigations of low-intensity ultrasound.

The finding of decreased firing during and after AM tFUS at 13 mW is one of the first reports of a reduction in spiking from tFUS. The majority of prior investigations have reported that tFUS increases firing in cortex (37; 52), hippocampus (8; 53; 54; 55), and others (56; 6). Note that reductions in evoked potentials, which have been reported with tFUS (11; 57; 58; 59; 60; 34; 61; 62; 63), do not necessarily imply a reduction in spiking. The amplitude of field potentials is strongly affected by the coherence among the recorded neurons (64), and thus it is difficult to relate changes in evoked potentials to underlying changes in firing rate. It is possible that the 40 Hz AM waveform employed here activated a distinct subset of ion channels (65; 66), or a different mechanism altogether than CW tFUS, leading to a net reduction in MUA activity. The finding that the reduction was not resolved at AM tFUS with a higher intensity (although it was evident when conditioning on the prevalence of SWR, see Fig 5C) suggests a complex interaction between acoustic intensity and waveform. It has been suggested that tuning the PRF may offer specificity of the neuromodulating effect from PW tFUS (37). Our findings indicate that both the timing of the ultrasonic perturbation and its force combine to shape the subsequent response to tFUS. Further investigation is needed to better understand the interplay between intensity and temporal dynamics, and how to optimally select their values to achieve the desired neurophysiological outcome. AM tFUS may provide a complementary tool in the tFUS arsenal by allowing for a transient reduction in activity.

The acoustic intensities employed in our study are well below the FDA safety guidelines for ultrasonic imaging: 720 mW/cm^2^. We were able to resolve firing rate changes at average intensities of only 13 mW/cm^2^ with AM tFUS and 52 mW/cm^2^ with CW tFUS. Furthermore, the mechanical index employed here (<0.02) is also an order of magnitude lower than the 0.3 suggested to be the upper limit of diagnostic imaging. Given the excellent safety profile of low-intensity ultrasound in imaging, it is very likely that the stimulation investigated here is safe. Moreover, recent studies that have tested tFUS intensities much higher than here (up to 25.8 W/cm^2^) reported no tissue damage as assessed by histological assessment of post-mortem brain tissue (67). The fact that neuromodulation was observed here at such low acoustic intensities is encouraging, as it implies that testing the effects of AM tFUS in humans may be carried out at power levels that are currently used in human ultrasound imaging practice, and thus unlikely to produce tissue damage. In addition to histological evaluation, temperature measurements, especially at the skull, will further lend evidence for the safety of AM tFUS.

Although our study provides evidence for the distinct nature of AM stimulation compared to CW tFUS, we did not perform a comparison between AM and PW tFUS. Future studies that match the PRF and AM frequency, as well as the average acoustic intensity, are thus required in order to ascertain the effect of the smoothness and continuity of AM waveforms on ultrasonic neuromodulation outcomes. There is evidence that abrupt pressure transitions may innervate the peripheral auditory system (15), with this side effect removed when smoothing the waveform edges. Another limitation of this study is the exclusive use of anesthetized animals. The state changes inherent to sleep and anesthesia are well-suited to investigating the gating of tFUS effects by baseline brain activity. Nevertheless, future studies are needed to understand the role of endogenous rhythms in the awake state on neuronal responses to focused ultrasound.

